# Self-organisation and positioning of bacterial protein clusters

**DOI:** 10.1101/133637

**Authors:** Seán M. Murray, Victor Sourjik

## Abstract

Many cellular processes require sub-cellular positioning of proteins. This can be due to passive mechanisms such as recruitment by existing landmarks or curvature sensing. However, in bacteria active self-positioning is likely to play a role in multiple processes, including the positioning of the future division site and cytoplasmic protein clusters. How can such dynamic clusters be formed and positioned? Here, we present a model for the self-organization and positioning of dynamic protein clusters into regularly repeating patterns based on a phase-locked Turing pattern. A single peak in the concentration is always positioned at mid-domain (mid-cell) while two peaks are positioned one at each quarter-position etc. Furthermore, domain growth results in peak-splitting and pattern doubling. We argue that the model may explain the regular positioning of the highly conserved Structural Maintenance of Chromosomes (SMC) complexes on the bacterial nucleoid and provides an attractive mechanism for the self-positioning of dynamic protein clusters in other systems.

## Introduction

Pattern formation in multi-cellular eukaryotic biology is often investigated using a reaction-diffusion mechanism in which a diffusion-driven instability results in the emergence of a pattern^1^. This was first applied to morphogenesis in a seminal paper by Turing^2^ and since then has become established in many systems in developmental biology such as skin pigmentation in fish, hair and feather follicle patterning in mouse and chicken embryos and the positioning of digit primordia within mouse limb buds^3,4^. Typical requirements for Turing pattern formation are a cooperative higher-order interaction^1^ and differing diffusion rates, or in the case of a membrane bound system^5^ or multi-cellular system^6^, differing reaction rates.

However, spatial organization is also essential within cells, where many cellular processes such as cell division, DNA segregation, cell polarity and motility require the positioning of proteins to specific locations within the cell. In bacteria, this is often achieved by passive mechanism such as recruitment or repulsion by existing landmark proteins or membrane curvature sensing^7^. Truly active mechanisms that do not rely on a pre-existing marker or gradient have largely been confined to DNA segregation. However, several systems have been discovered that appear to exhibit active positioning, especially, but not only, in the context of choosing the future division site^8,9^. How can protein complexes, which can be very dynamic and exhibit rapid turnover, be positioned? Geometry sensing can occur due to a difference in the local bulk volume and unequal membrane affinities^10^. However, the resulting pattern labels only geometrical distinct regions and cannot produce a repeating regular pattern along the length of a rod-shaped cell.

Turing-type mechanisms on the other hand, have not been applied very much in the intra-cellular context (see however ref.^11^ for the case of yeast polarity). This is probably due to the sensitivity of the mechanism to initial conditions^12–14^ and parameters^15^ – the ‘fine-tuning’ problem of mode selection and the ‘robustness problem’ of maintaining a given pattern^16^. The pattern, when it exists, does not generally have a fixed pre-determined phase. Fixed (Dirchelet) boundary conditions can help in pattern selection^12,17^ but are not relevant in the subcellular setting where zero-flux (Neumann) boundary conditions are more appropriate (certainly for systems not relying on pre-existing pattern). Noe that while classically referring only to static patterns (the focus of this work), the Turing mechanism can also induce oscillatory patterns, which have been studied extensively in the context of the Min system (see ref.^18^ for a review).

In this work, we present a model for the self-organization and positioning of dynamic protein complexes using a phase-locked Turing pattern. The model produces a regular repeating pattern that is insensitive to initial conditions and has a fixed phase. Nucleoid-cytosol exchange essentially selects and phase-locks the Turing pattern even though many modes are linearly unstable, overcoming the drawbacks mentioned above. In short cells the mechanism results in a single focus at mid-cell, while in longer cells a focus is positioned at each quarter-position and so on with increasing length. Furthermore, the system exhibits pattern doubling due to domain growth and can be controlled and made more precise by inhomogeneous nucleoid binding and unbinding. Importantly, the mechanism does not rely on any geometrical input other than a bounded domain. The model is motivated by the self-positioning of the Structural Maintenance of Chromosomes (SMC) protein complexes^19^ on the bacterial nucleoid but is generally applicable to other systems.

## Results

### A model of MukBEF cluster formation and positioning

Ubiquitous in all domains of life, Structural Maintenance of Chromosomes (SMC) protein complexes are condensins, required for correct chromosome condensation, organization and segregation. MukBEF, the *E. coli* SMC, consists of a dimer of MukB, joined at a hinge domain and at an ATP-binding head domain to form a loop capable of entrapping DNA, together with the small accessory proteins MukEF, which interact with MukB in an ATP-dependent manner (see Rybenkov et al.^20^ for a review). The consensus from *in vivo* and *in vitro* studies is that ATP serves as a DNA-binding switch: ATP binding promotes loading onto DNA, while ATP hydrolysis stimulates detachment. Studies have shown that MukBEF forms clusters in the middle of the nucleoid in short *E. coli* cells^21–24^ and this localization changes to the two quarter positions in longer cells. However, the mechanism of this positioning is unknown. A live-cell study has indicated that the fluorescent foci consist of relatively immobile complexes, while another pool diffuses much more rapidly^25^. The immobile foci were found to be composed of 8-10 MukBEF dimer of dimers complexes that were suggested to have entrapped multiple DNA strands^25^.

How do MukBEF clusters form and how are they positioned? Non-specific DNA binding and the requirement of ATP hydrolysis^26^ suggest a directed energy-dependent mechanism, rather than simple recruitment. We hypothesized that the system could be described by Turing pattern formation. This was motivated by two properties described above: 1) two populations with differing diffusion constants and 2) the presence of cooperative (higher-order) reactions.

The general model scheme consists of two ‘species’ *u* and *v* interacting in a bounded one-dimensional ‘bulk’ over a one dimensional surface (Fig. 1a, upper panel). Species *u* exists in the bulk (the cytosol) with concentration *u*_*c*_ and on the surface (the nucleoid) with concentration *u*_*m*_. Species *v* exists only on the surface with concentration *v*_*m*_. Species *v* diffuses slower than *u*. In the context of MukBEF, *u*_*m*_ represents the concentration of the free form of the basic functional complex, the dimer of dimers, while *v*_*m*_ is the concentration of the slower diffusing form of the complex, due to entrapment of DNA. The former becomes the latter at a basal rate α and cooperatively at a rate *β*, and the latter becomes the former at a rate γ. We envisage the cooperative conversion to be due to transient interactions aiding the engulfment of DNA by each of the two constituent dimer rings. In this way, slowly-diffusing complexes cooperatively recruit quickly-diffusing subunits. The concentration of the cytosolic state is given by *u*_*c*_, which converts to and from the bound states with linear rates ϵ and, δ and δ’ respectively. Figures 1a, b show a diagram of the proposed model and the system of partial differential equations describing it. For reasons of computational complexity, we largely restrict ourselves to one spatial dimension. This is justified since *E. coli* is a rod-shaped bacterium and by the large size of fluorescent foci in comparison to the nucleoid^25^. With the parameter set used, the system has a single fixed point (Fig. S1a).

**Figure 1.**
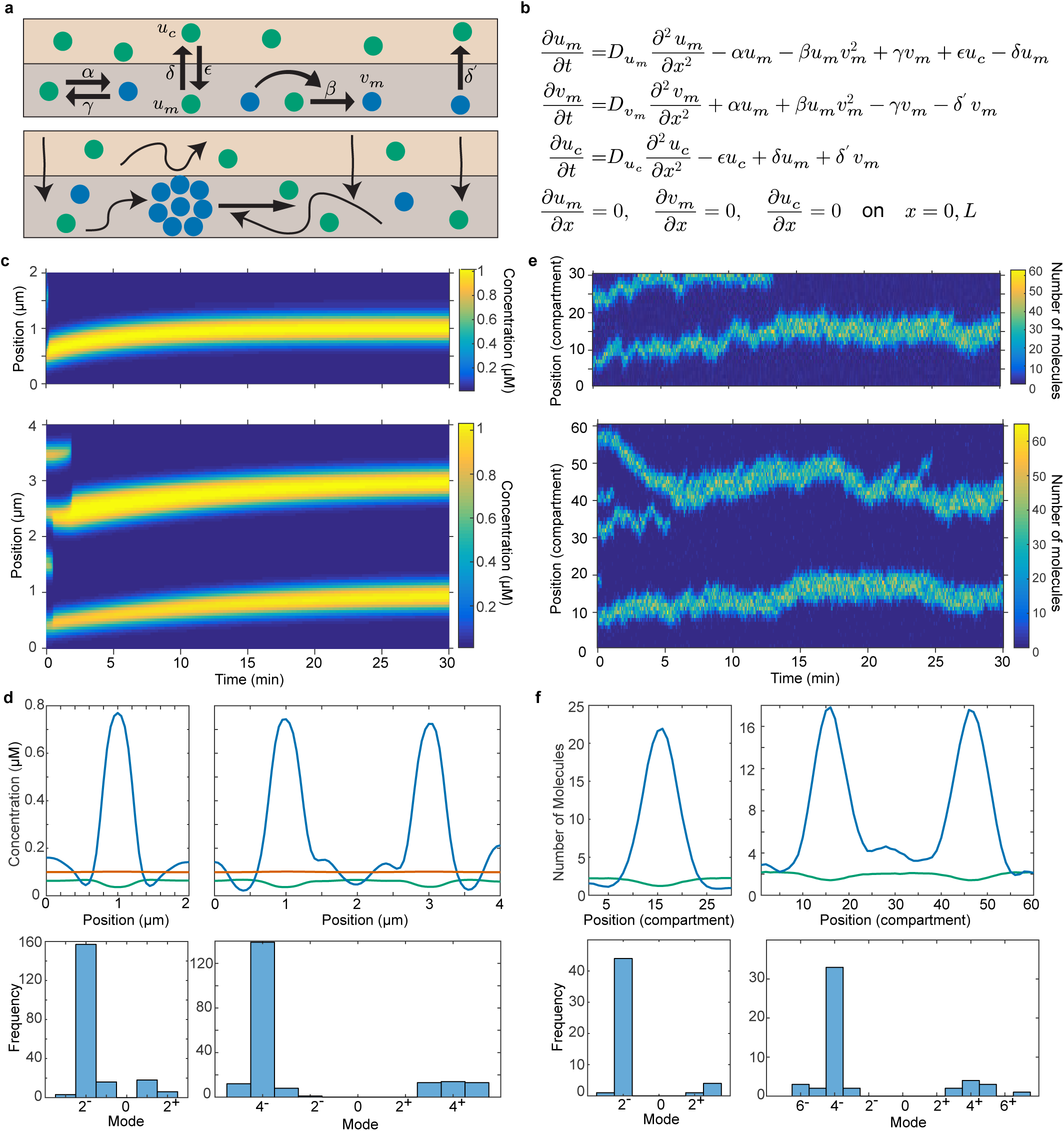
A self-positioning Turing pattern. (a) (Upper panel) Schematic showing the reactions of the system (see text). Species *u* (green) exists in the bulk (lighter color) or on the surface (darker color). Species *v* exists only on the surface. Binding and species interaction are indicated by arrows. Diffusion is not shown. (Lower panel) Schematic showing the flux balance mechanism. Thinner arrows represent binding or diffusion. Thick arrow indicates the direction of cluster movement. (b) The partial differential equations describing the system. *L* is the domain (system) length. (c) Kymographs of *v*_*m*_ from a solution of the equations in (b) over 2 and 4 μm domains. (d) The system was solved for 200 different random initial conditions. (Top) The average distribution of *u*_*m*_ (green), *v*_*m*_ (blue) and *u*_*c*_ (red) after 30 min are shown for 2 μm and 4 μm domains. The contribution of the boundary modes are visible. Note the homogeneous cytosolic distribution (red). (Bottom) Histograms of the dominate non-zero mode at the end of each simulation. See Fig. S2b for the case of a 3 μm domain. (e) Example kymographs of *v*_*m*_from stochastic simulations over 3 μm and 6 μm domains. See Fig. S5 for examples of single long-time simulations. (f) The stochastic system was ran 200 times as in (e) and the distributions (top) of molecules of *u*_*m*_ (green) and *v*_*m*_ (blue) were obtained by averaging over the last 10 min of the simulations. Boundary modes are not visible above stochastic effects. (Bottom) Histograms of the dominant mode of each simulation.

### A self-positioning Turing pattern

On solving the equations in Figure 1b numerically, we found that the system can self-organize with localized high concentrations of species *v* (Fig. 1c). Interestingly, we observed that the phase of the pattern is nearly always fixed. For our choice of parameters, there was generally a single peak in *v*_*m*_ positioned at the midpoint (mode 2^-^, see methods) on a 2 μm domain, whereas there were peaks at each quarter position (mode 4^-^) on a 4μm domain (Figure 1c, d). This is stark contrast to the prediction from linear stability analysis (Fig. S1b) that many modes, both even and odd, are driven spatially unstable by diffusion with modes, n=4 and n=8 respectively, having the greatest growth rates (see methods for further details).

For a small set of initial conditions, foci stabilized on the boundary (odd modes or even modes with positive amplitude), breaking the phase fixing (Fig. 1d, Fig. S2a). However, for all initial conditions that resulted in no boundary foci, the phase of the pattern was fixed as above. Furthermore, this was the case even for initial configurations far from the homogeneous steady state. Note that the Turing mechanism employed here does not produce static foci, rather the peaks constitute regions of high molecular density, with molecules free to diffuse in and out of the region – the model contains no oligomerisation beyond that of the basic functional complexes. We also solved the system in three dimensions and found similar behaviour (Fig. S3).

To better understand the nature of this pattern fixing, we considered the system without the bulk form of *u* - both *u* and *v* remain confined to the surface. In this reduced system, very similar patterns formed but they were not self-positioned. Foci formed within a few seconds but did not move even over very long timescales (Fig. S4a, b), though they could, as in the full system, disperse before the final pattern stabilized. Stable patterns were dominated by a broader distribution of modes though the mid-domain (mode 2^-^) was the most likely foci location (Fig. S4c). This was also true over a 4 μm domain, which generally has foci at the quarter positions in the full system (Fig. 1d).

These results demonstrate that binding and unbinding of molecules is necessary for dynamic self-positioning of foci. We explain this effect by a flux-balance argument (Fig. 1a, lower panel). The flux (number per second) of *u* molecules binding from the bulk to any particular region is proportional to the region area. Now suppose a high density focus of *v* molecules has formed off-center. Then, the flux of *u* molecules reaching the focus from each side is proportional to the area (length) of the domain on each side. The difference in flux from either side results in a locally higher reaction rate from *u* to *v* on one side than the other, which, in the case of a single focus, causes the focus to move to mid-cell, where the fluxes on either side balance. On a larger domain, where there is more than one focus, the same argument leads to regular positioning. See the SI text for further discussion. This very general argument has been previously made for regular plasmid positioning^27^ and, albeit implicitly, for positioning of the lateral chemotactic clusters in *E. coli*^28^, which are both essentially static objects. Note that only the former is an example of self-organisation^29^; the latter system exhibits self-assembly ^30^. Here, however, nucleoid-cytosol exchange, in combination with a Turing instability, results in the self-organisation and positioning of highly dynamic protein clusters. Importantly, unlike the plasmid case, the positioning is done by the same proteins that constitute the object, rather than a separate positioning system.

### Stochastic effects increase robustness of positioning

Biological systems are subject to noisy environments and this is especially true for proteins with small cellular concentrations. Indeed many (potential) self-positioning proteins have relatively small molecular numbers and therefore their positioning is continually subject to perturbing stochastic effects. Recent estimates for the number of MukBEF molecules in the cell are in the range of 200-500 molecules^25,31^. We therefore developed a stochastic (Gillespie) simulation of the system to investigate the effect of noise on the positioning mechanism.

We again found pattern-fixing behaviour. Foci fluctuated around the middle position (3 μm domain) or at the quarter positions (6μm domain) (Fig. 1e). The distributions obtained over long-time simulations showed clear maxima at these positions (Fig. S5). Furthermore, averaging many independent simulations reproduced this distribution (Fig. S2c). As for the deterministic case, foci occasionally developed on the domain boundaries but were much rarer (Fig. 1f). Most importantly such asymmetric patterns were not stable and after a short time the system returns to the correct position (Fig. 1e, Fig. S5). We also found that positioning is lost in the absence of exchange with the bulk (Fig. S4d, e). The flux balance mechanism evidently biases selection of both the desired regularly positioned symmetric modes and the undesirable boundary modes, while stochastic effects destabilize the latter (see methods and SI Text). Furthermore pattern formation and positioning occurred over a relatively wide parameter range, especially in the stochastic case (Fig. S1c, methods). These results demonstrate the robustness of the patterning and its independence on initial conditions.

### Peak-splitting is induced by growth

Positioning mechanisms in bacteria are often involved in the partitioning of chromosomes and plasmids. As such, the timely splitting and re-positioning of clusters is usually central to their function. We therefore incorporated domain growth into the simulations in order to examine the effect on patterning and whether it is sufficient to induce splitting. When we considered a single focus on an exponentially growing domain, we indeed observed peak-splitting where the mode 2^-^ solution abruptly splits into the mode 4^-^ solution (Fig. 2a). Treating the domain length as a bifurcation parameter, we found the critical domain length to be L_crit_=5.45 μm (Fig. 2b). Interestingly, we observed that a shrinking domain causes the opposite bifurcation but at a lower critical length of 2.71μm, indicating a type of hysteresis (Fig 2b, Fig. S6a). This bifurcation does not occur via peak merging, rather one of the two peaks disperses, while the other moved to the mid-domain position. We found that the critical length L_crit_ increases with both on-nucleoid diffusion rates. However, increasing the cytosolic diffusion constant has no effect since the cytosol is already well-mixed on the timescale of protein (un-)binding (SI Text).

**Figure 2.**
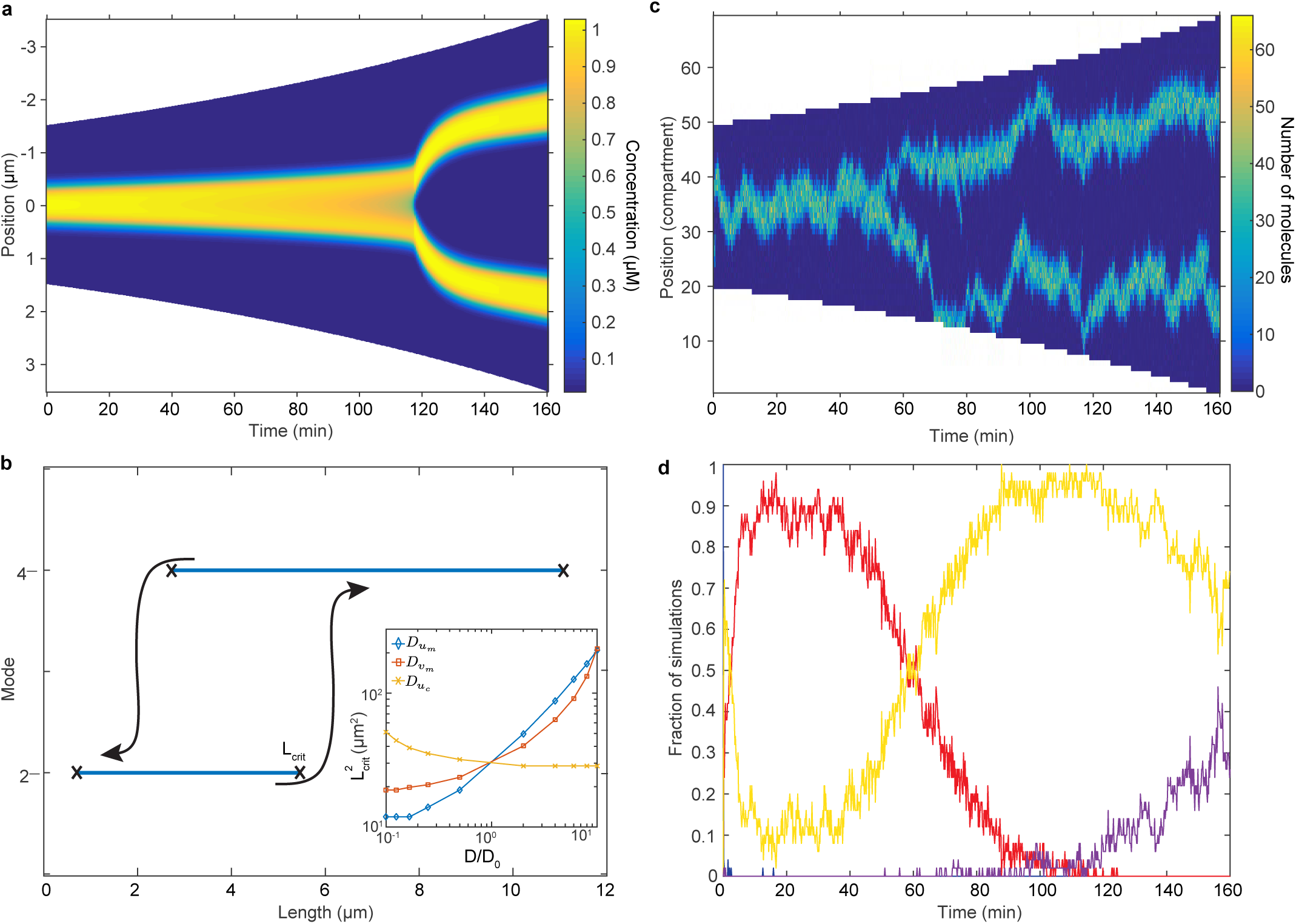
Pattern splitting during exponential domain growth. (a) A kymograph of *v*_*m*_ from the deterministic system during exponential growth starting from the mode 2^-^ solution (doubling time is 130 min). See Fig. S6a for domain shrinking. (b) Diagram indicating the domain lengths for which the deterministic 2^-^ and 4^-^ patterns are stable and the bifurcation points. Arrows indicate hysteresis-like behaviour. (Inset) The relationship between the L_crit_^2^ and the diffusion constants of the model. (c) Example kymograph of *v*_*m*_ from a stochastic simulation with exponential growth (doubling time as in (a)). The initial domain length is 3 μm. See Fig. 3a for an average over many simulations. (d) The fraction of 50 stochastic simulations with 0 (blue), 1 (red), 2 (yellow) or 3 (purple) peaks as a function of time. The transition from one to two peaks occurs within an approximately 40 min window. See Fig. S6b for the average number of foci as a function of time.

Turning to stochastic simulations, we found much the same behaviour with one focus at mid-domain rapidly becoming two foci at the quarter positions (Fig. 2c). Peak-splitting occurred earlier than in the deterministic simulations and generally between 40 and 80 mins into the simulation with half of the simulations having split by 60 min / 4.1 μm (Fig 2d), whereas the critical length L_crit_=5.45 μm is reached later at 112 min. It therefore appears that the bifurcation here is somewhat analogous to a saddle-mode bifurcation in that the basin of attraction of the stable fixed points shrinks as the bifurcation is approached. Stochastic fluctuations then allow the system to jump to the mode 4-branch before the bifurcation occurs.

We next investigated extended domain growth. It has already been shown that exponential domain growth can lead to robust pattern doubling of Turing patterns in the deterministic case^32^ and we found this to also be the case here (Fig. S7). On the other hand, pattern doubling has been found not to be robust in the stochastic case^13^. Indeed, we observed that new foci appear as the domain grows and are regularly positioned but they do not split simultaneously like in the deterministic case (Fig. S8a). However, we found that the average number of foci as a function of the domain length shows a clear linear relationship with 1 additional focus every 3 μm (or, equivalently, an exponential relationship with time) (Fig. S8b). Therefore, while foci do not split synchronously in individual simulations, the average growth behaviour shows pattern doubling. Similar regular positioning of foci is found in *Streptomyces coelicolor*^33^. During sporulation, the protein SsgA forms many regularly positioned foci within (multi-chromosome) long aerial hyphae and serves to positively regulate the position of the future cell division site.

In order to qualitatively compare these results on peak splitting with the biological system, we imaged a strain carrying a MukB-GFP fluorescent fusion at the original chromosomal location^25^. As mentioned above, MukB forms clusters at the mid- or quarter-cell positions. Furthermore, these clusters can be composed of one to two closely spaced foci, which interconvert reversibly and dynamically within a timeframe of less than 5 min^34^, reminiscent of what we observe in our individual simulations (Fig. 1e, Fig. 2c, Fig. S5, Fig. S8). This behaviour made it difficult to ascertain when precisely a cluster had irreversible split using time-lapse experiments (Fig. S9a-c) especially when we wanted to relate this to the time at which the nucleoid becomes bi-lobed (see below), an event which is also initially dynamic and reversible. Furthermore, photo-bleaching restricted the frame rate of time-lapse experiments to no faster than a frame every 3 min. We therefore took a demographic approach^35^, imaging many cells with high image quality and extracting the MukB-GFP fluorescent profile along the long cell axis. Profiles from cells having the same length, to the nearest two pixels (130 nm) were then averaged and arranged into an averaged demograph (Fig. 3b). We found a pattern very similar to that of average over many stochastic simulations (Fig. 3a) with a clear relationship between the cell length and the number of MukB-GFP clusters. While these experimental observations do not directly prove that splitting of MukB-GFP clusters is due to growth, they are qualitatively consistent with the growth induced splitting in our model.

**Figure 3.**
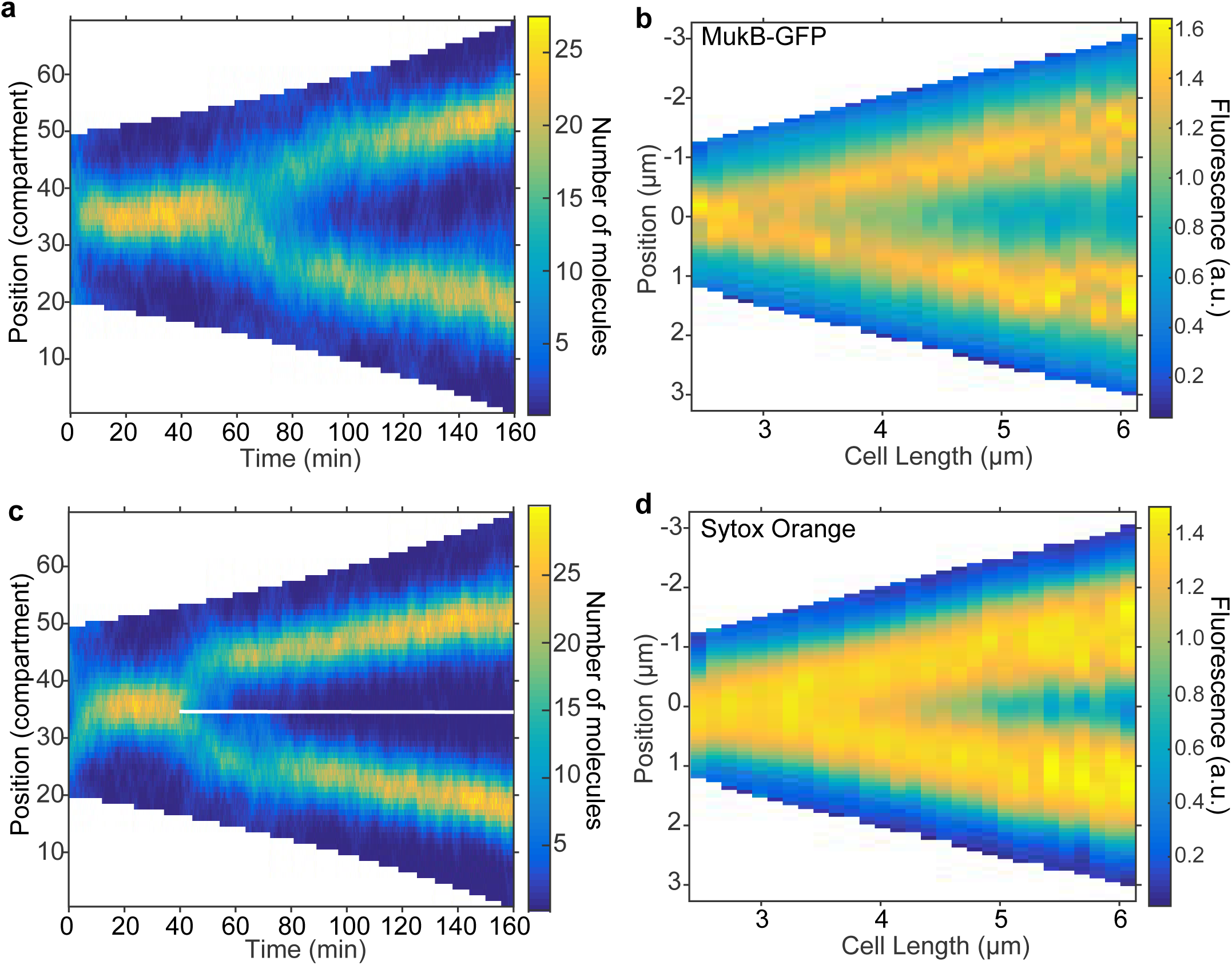
The model provides an explanation for MukBEF clustering and splitting. (a) An average kymograph of *v*_*m*_ from 50 stochastic simulations as in Fig. 2a. (b) An averaged demograph from 884 cells showing the MukB-GFP distribution as a function of cell length. (c) As in (a) but the surface is divided into two from 40min (white line). Exchange between the two halves occurs via the bulk. (d) An averaged demograph from the same cells as in (b) showing the nucleoid distribution (stained with Sytox Orange) as a function of cell length. See also Fig. S9. In (a) and (c) the total concentration of molecules is constant. Note the different x-axis between simulated kymographs and experimental demographs. Model parameters are the same as in the static case and have not been specifically chosen to obtain agreement between (a) and (b).

### Nucleoid compartmentalization can enhance peak-splitting

Nucleoid-bound proteins are inherently affected by chromosome segregation: diffusion between the two nucleoid lobes is likely to be severely restricted with the majority of protein exchange occurring via the cytoplasm. To investigate how this affects pattern splitting, we mimicked chromosome segregation in our simulations by splitting the surface (the nucleoid) into two compartments, while still allowing exchange via the bulk (the cytoplasm). We performed the simulations as previously but now with a compartmentalized surface from 40 min onwards. Interestingly, we found this was able to induce focus splitting earlier and more synchronously than domain growth alone (Fig. 3c, Fig. S6b). This suggests that chromosome segregation can play a role in the positioning and splitting of self-organized protein clusters. This may especially be the case in systems that have many repeating foci such as SsgA. Compartmentalization via chromosome segregation then essentially reduces the problem of forming and positioning multiple separated foci to simply forming and positioning multiple single foci individually and may therefore contribute to the robustness of these systems.

Does nucleoid segregation play a role in the splitting of MukBEF clusters? To address this question, we used a DNA stain, Sytox Orange, to accurately stain the nucleoid in live cells and determine whether nucleoid segregation (as measured by a bi-lobed nucleoid) begins before or after MukB-GFP clusters split. Time lapse experiments indicated that these two events are reasonable synchronous but as discussed above their dynamic nature and the restricted frame rate made it difficult to draw any further conclusion. We therefore performed a demographic analysis as for MukB-GFP (Fig. 3d). We found that the nucleoid starts to become bi-lobed only after the MukB-GFP cluster has split (Fig. S9f). This suggests that nucleoid segregation is not the cause of cluster splitting. While the effect we observed was subtle (but statistically significant) and based on average demographs, two previous observations support the conclusion. Firstly, depletion of the topoisomerase TopoIV, an enzyme that is essential for chromosome separation, produces elongated cells with several catenated chromosomes and a homogeneous nucleoid but several evenly spaced MukB foci^36^. Secondly, the nucleoid becomes bi-lobed from about half-way through DNA replication^37,38^ whereas MukB foci splitting is approximately coincident with *ori* separation^34^, an earlier event. The growth induced splitting of our model therefore provides an attractive alternative mechanism for MukBEF cluster splitting.

### Inhomogeneous binding and unbinding of proteins modulates peak positioning

In the model presented thus far, we have taken binding and unbinding of proteins to be uniform. However, many nucleoid associated proteins exhibit some amount of specific DNA binding. Both *E. coli* MukBEF and *B. subtilis* SMC do not exhibit sequence specific DNA binding^26,39,40^. However, the presence of other proteins cause the latter to be specifically loaded onto the DNA at sites in the *ori* region^41,39^, while the former is specifically unloaded from the DNA at sites in the *ter* region^26^.

We therefore incorporated site-specific loading and unloading into the model. We found that a specific binding site at a quarter position had an attractive effect on positioning, pulling the focus away from the mid-domain position (Fig. S10a). Interestingly, we observed that two distant higher-affinity sites do not result in the formation of two foci over a domain length (3 μm) that only supports a single focus in the homogeneous case (Fig. 1f).

A specific unbinding site had the opposite effect, pushing the focus towards the middle of the larger side of the domain and away from the centre position (Fig. S10b). At high unbinding rates, the site effectively splits the domain into two, similar to a diffusive block. However, there was almost always one focus present at a time, evidently since the domain size (3 μm) can support only focus (Fig. 1f). In summary, inhomogeneous binding or unbinding can affect focus positioning but not induce splitting.

### Inhomogeneous binding can enhance peak splitting

We already observed that focus splitting can occur solely due to domain growth and therefore wondered what effect inhomogeneous binding and unbinding would have in the presence of growth. We found that maintaining a single specific binding site at the mid-domain position with a binding rate 15 times that of elsewhere was able to delay focus splitting for approximately 60 min, almost half the doubling time (Fig. 4a, d). While a single focus was present, this resulted in more precise positioning and a higher local concentration of molecules and the opposite while there were two foci (compare with Fig. 3a, S6b). The factor of 15 was chosen based on Fig. S10a so that the value is below the saturating regime. Interestingly, the effect on the time of splitting was not as strong when we maintained a higher-affinity site at each quarter position (Fig. 4b, e). In that case splitting occurred around 10-15 min earlier. However, the positioning was more precise while there were two foci and less precise while there was a single focus. The latter effect was due to the single focus stochastically switching between the neighbourhoods of the two higher-affinity sites. Finally, we considered a specific unbinding site maintained at the mid-domain position. We found an approximately 15 min delay in focus splitting and mis-positioning when a single focus was present (Fig. S10c, d). The focus generally stabilized on one side of the mid-position, resulting in a bi-modal distribution on average.

**Figure 4.**
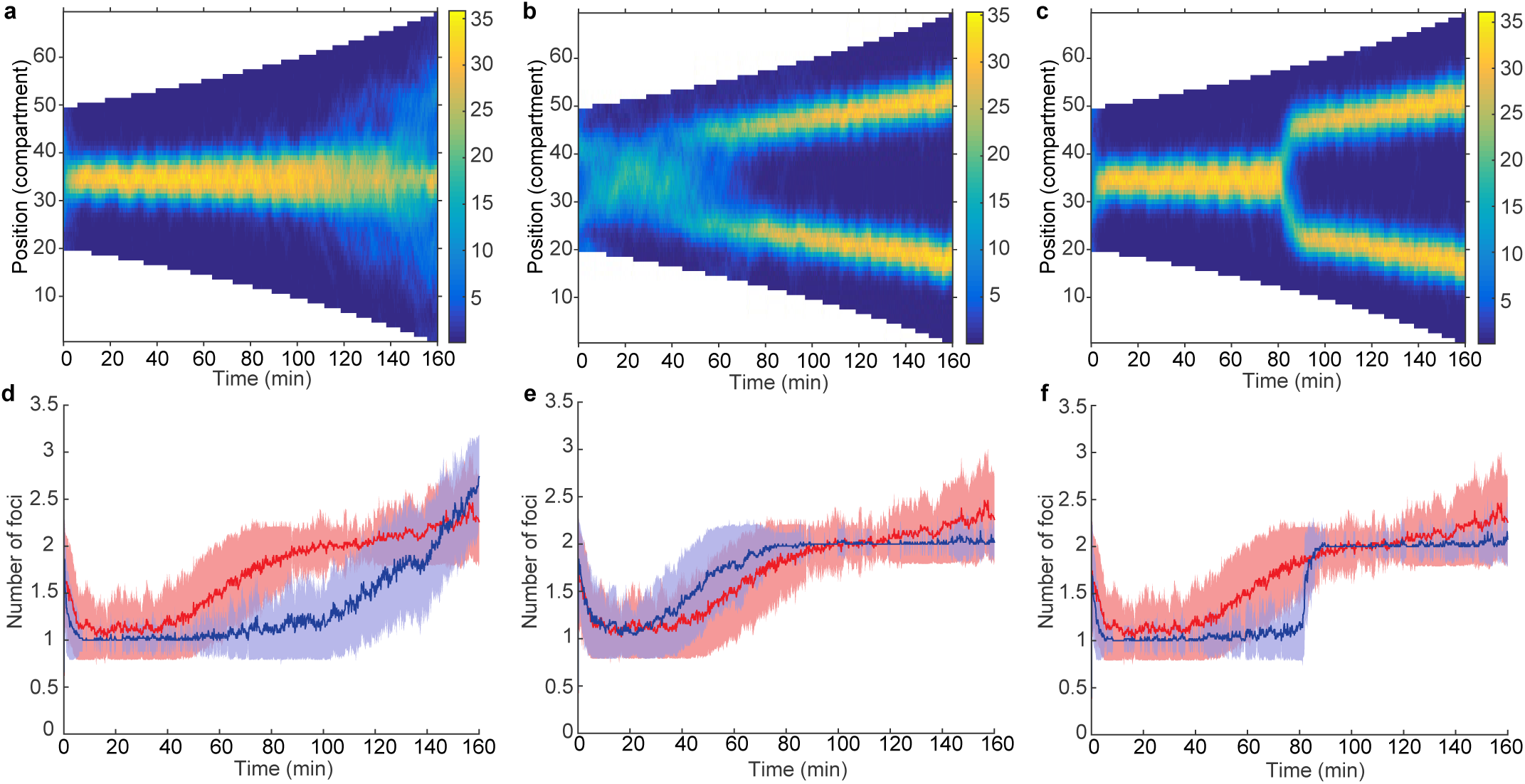
Effects of inhomogeneous binding during exponential growth. (a-c) Average kymographs of *v*_*m*_ from 50 stochastic simulations for the cases of a single specific binding site at mid-domain (a), two sites at the quarter positions (b) and of a single site at mid-domain that splits into two sites, one at each quarter position at 80 min (c). The corresponding mean number of foci is shown in blue in (d), (e) and (f) respectively. For comparison, the homogenous case from Fig. 3a and Fig. S6b is shown in red. The shaded region represents one standard deviation. The specific binding rate is 15 times that of the other compartments. Simulations were otherwise performed as in Fig. 3a.

Every chromosomal site is duplicated and positioned during the cell cycle. For example, the *ori* regions in slow-growing *E. coli* cells are positioned at mid-cell before being replicated and migrating rapidly to the quarter positions^37,42–45^. We therefore asked what effect a specific binding site following such a pattern would have on focus position and number. For simplicity, we simply implemented a discrete duplication and positioning event 80 min into the simulation i.e. a single higher-affinity site at mid-domain turns into a site at each quarter position. At this time point in the homogeneous case, 80% of simulations had already split their focus (Fig. 2d). However, we found that site duplication was able to accurately set the time of splitting to the 80 min time point (Fig. 4c, f). It only required about 10 min for all the simulations to split and reposition their foci. Furthermore, as the ‘default’ positions of the foci matched the locations of these sites, the localization was more precise throughout the entire simulation than in the case of homogeneous binding.

These results indicate that while specific binding (or unbinding) is not necessary for the self-positioning mechanism described here, it can have interesting effects on patterning. When site-specific binding is strong such sites can override the inherent positioning of the mechanism, while at lower, perhaps more realistic, strengths the effect is mildly attractive. When the location of site-specific binding site coincides with the site ‘preferred’ by the mechanism then this additional positive influence results in more precise positioning. And when this is combined with duplication and segregation of the specific binding site, foci-splitting is also made synchronously.

## Discussion

### MukBEF localization as a self-positioning Turing pattern

The *E. coli* SMC complex, MukBEF, is required for efficient chromosome segregation and organization. Through an unknown mechanism, it forms nucleoid-bound clusters at mid-cell or, in longer cells, at the quarter positions. Here we have proposed an explanation for this behaviour in the form of novel self-positioning Turing-type mechanism. We developed a simple mathematical model based on the current understanding of the system and used measured diffusion and binding rates. We found that the system is capable of self-organization, forming high-density regions (foci) spontaneously and in the absence of stable complex-complex interactions. Furthermore, the exchange of molecules between the nucleoid and the cytosol results, via balancing fluxes, in foci positioning, placing a single focus at mid-cell or, in longer cells, a focus at each quarter position. We found unintuitively that stochasticity improved the accuracy of positioning in that sometimes foci stabilised on the boundary in the deterministic case.

Two observations further support our model of dynamic self-positioning. Firstly, positioning is perturbed in a strain with a mutant MukBEF that can bind but only weakly hydrolyze ATP^26^. Since ATP release results in unbinding from DNA, this mutant is described in our model by blocking unbinding from DNA. When we did this, we also found mis-positioned foci (Fig. S4). Secondly, repletion of the MukF sub-unit in non-replicating cells (which generally have a single focus at mid-cell), leads first to the appearance of multiple weak MukB-GFP foci distributed across the nucleoid that then take up the wild-type number and position over time^34^. This is consistent with what we observe in our simulations, where we start with a homogeneous concentration of molecules as in the repletion experiment (Fig. 1e).

Recent work has uncovered further complexity in this system. In the aforementioned ATP-hydrolysis mutant strain MukBEF foci actually associate with *ter* rather than *ori*^26^. It is believed that MatP-*matS* promotes unloading of MukBEF by stimulating ATP hydrolysis. However, it is still unclear what underlies the association of MukBEF to *ori* and, in *ΔmatP* cells, to *ter*. In our simulations, we showed that sites with a higher MukBEF loading rate have an attractive effect on foci positioning, whereas sites having a higher unloading rate had a repulsive effect. The former is consistent with the association to *ori* and, in the absence of MatP, to *ter*, while the latter is consistent with the lack of *ter* associated foci in the presence of MatP. There is also evidence that MukBEF foci play a role in positioning segregated origin regions, recruiting them to the mid-cell or quarter positions^24,36^. This intriguing possibility is now bolstered by the self-positioning mechanism that we have presented here.

### Self-organisation and positioning of dynamic protein clusters

Protein clusters can be very dynamic with rapid turnover and yet be regularly positioned within the cell. The model presented here explains this behaviour using a Turing-type reaction diffusion mechanism for self-organisation. Positioning is due to surface-bulk (nucleoid-cytosol) exchange leading to a disparity in fluxes on the surface. A similar flux-balance argument has been proposed for the positioning of (non-dynamic) plasmids by the ParABS system^27^. Here, however the foci (protein clusters) are dynamic objects that self-organize and can position themselves independently without relying on any external positioning proteins. Furthermore, while flux-balance can provide the spatial information to position plasmids, the actual molecular mechanism for plasmid movement is unknown, with several competing models^27,46,47^. Here, however, the self-organisation and positioning are self-contained, requiring only the basic reactions of the system (Fig. 1a).

The key ingredients of the model are the following: a protein with cooperative self-interactions that exists in two nucleoid-bound populations with differing diffusion rates and for the positioning mechanism, exchange of molecules with the cytosol. Additionally, energy expenditure (e.g. ATP hydrolysis) is required for self-organization^29^ and the positioning mechanism requires that this be linked to unbinding from the nucleoid. Importantly, oligomerisation is not a requirement.

Given the generality of the model, we expect it to applicable to other systems. For example, the cytoplasmic chemotactic clusters in *Rhodobacter sphaeroides*^48^ and the PomZ cluster of *Myxococcus xanthus*^49^ both exhibit regular positioning and require non-specific DNA binding and ATPase activity.

### A phase-locked Turing pattern

Finally, from a mathematical viewpoint, this work presents a reaction diffusion system that results in a specific pattern with a fixed phase that is not sensitive to initial conditions and is robust to perturbations in parameters. Such phase-fixing has typically been achieved by tuning the model parameters such that a single mode is driven unstable by diffusion and hence dominates the final pattern^1,12^ or by using fixed or mixed boundary conditions^1,12,17^. Here, however, the patterns produced are far from equilibrium and are similar in that regard to the spike solutions found analytically in other systems in the limit of asymptotically large diffusivity ratios^50^. Here, the solution peaks are broader and are composed of just a few modes. Like spike solutions their shape is not sensitive to the initial perturbation that created them. However, we showed that stochastic effects cause the ‘undesirable’ boundary patterns to be destabilised, ensuring that *which* pattern is formed is also independent of the initial perturbation i.e. the stochastic pattern consists solely, on average, of a parameter and domain-size dependent number of interior regularly positioned peaks. This result will therefore contribute to the application of the Turing mechanism in explaining biological pattern formation.

## Materials and Methods

### Deterministic simulations

The system of partial differential equations was solved in Matlab using the pdepe solver. The system was integrated over a mesh with a spacing of 0.05 μm. Initial conditions were taken to be a random perturbation about the homogeneous steady state (drawn from a normal distribution with standard deviation of 10%). For simulations involving growth, the Lagrangian formalism was used for computational efficiency and the solutions mapped back to the Eulerian formalism. The two dimensional solution was obtained using a custom-written code based on the method of finite differences.

### Stochastic Simulations

Exact stochastic (Gillespie) simulations^51^ were implemented in C++ and based on the Enhanced Direct Method (EDM) of Gibson and Bruck^52^. We also combined the sorted linear ordering of propensities of the Optimized Direct Method (ODM)^53^ with the binary tree search of the EDM: The propensities were sorted into reaction types (Diffusion of species 1, Diffusion of Species 2, Binding of species 1 etc.) and stored with their partial sums in individual binary trees. The roots of these trees were then sorted in descending order to be searched linearly as in the ODM, followed by a binary search up the tree as in the EDM. We used the Ziggurat Method for generating random variables. In the stochastic simulations the bulk is assumed to be well-mixed. This is justified due to the much slower timescale of binding/unbinding compared to that of diffusion, which results in a homogenous bulk concentration (Fig. 1d, red line) and by the independence of one to peak transition on the higher cytoplasmic diffusion rates (Fig. 2b). The spatial aspect is dealt with by dividing the domain into compartments, each having a width of 0.1 μm (apart from the simulations in Fig. S8, which used 0.15 μm) and between which the species can diffuse. Initial concentrations were set to the integer homogenous configuration closest to the continuous model. The stochastic system was simulated using the same fundamental parameters as for the deterministic system (appropriately converted from units of nM and μm to number of molecules and compartment length respectively). The system state was read out every 5 or 10 s. For simulations with growth, the simulation was paused after every time-duration that corresponded to growth by one compartment. An additional (empty) compartment was then inserted at a random position and the volume and total number of molecules (via the cytosolic fraction) were increased, maintaining the same overall concentration.

Note that the domain lengths over which either one or two foci were consistently found are different in the two simulation methods due to stochastic effects. The stochastic case requires longer domains (3μm and 6μm respectively) than the deterministic case (2μm and 4μm) for the same parameter set (compare Fig. 1d, f and S2b). Intermediate domain lengths can have one or two foci depending on the initial conditions in the continuous case or stochastic switching between one or two positioned foci in the stochastic case. The domain lengths used are chosen to differ by a factor of two as we also discuss domain growth (doubling), which has the effect of locking in a particular wave number k.

### Pattern analysis

Linear stability predicts that the solution around the fixed point is rendered spatially unstable due to the presence of diffusion and is composed of a discrete set of fundamental modes of the form cos(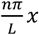). The integer n as the mode number (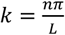 is the wave number) and we denote the sign of the amplitude of this mode with a superscript e.g. mode 2^-^ has a peak a mid-domain, whereas mode 2^+^ has a valley at mid-domain (peaks on the boundary). Note that only even modes with negative amplitude correspond to interior (regularly positioned, symmetric) peaks. We refer to odd modes and even modes with positive amplitude as boundary modes. The growth rate of each mode around the homogeneous fixed point is given by the dispersion relation and the fastest growing unstable mode is expected to dominate the final pattern^1^. However, higher order perturbations and multiple unstable modes can result in this prediction not holding^1^. For the chosen parameters over a 2 μm domain, linear stability would predict that the mode n=4 (wave number *k*_*m*_ = 6.5) will dominate in both the reduced and full models (Fig. S1b, red line). However, we find that the mode 2^-^ pattern is selected, robustly in the stochastic case (Fig. 1d, f). In Fig. S1b, the first 10 (relative) amplitudes of the mode expansion are shown for some example patterns over 2 μm (deterministic cases) and 3 μm (stochastic case) domains. The patterns of the full deterministic and stochastic models are largely composed of only even modes with the largest contribution coming from mode 2^-^ (see also Fig. 1d, f).

Evidently, the presence of higher-order perturbations and/or the range of unstable modes mean that the linear approach cannot be used to predict the long-term behaviour of the system. Indeed higher order effects are supported by the observation that at the very beginning of the simulation, the pattern is often consistent with the predicted mode and before subsequently changing. A mathematical analysis taking these higher order effects into account is beyond the scope of this work but we note that some progress has been made in that direction^54^. Non-linear analyses have also been used to study far-from-equilibrium solutions of reaction-diffusion equations. For example, spike or pulsed solutions have been studied extensively in the limit of asymptotically large diffusion ratio (see^50^ for a review).

### Reduced Model

In the reduced model, we set all binding and unbinding rates to zero and started the (two-variable) simulations with the same number of bound species as in the full system. This can equivalently be seen as taking the limit ϵ, δ → 0, while keeping ϵ/(ϵ+δ)=0.75. Furthermore, it is not difficult to see that since γ≫ Δ, the fixed points of the two systems are almost identical. As described in the text, correct positioning was no longer obtained in either the deterministic or stochastic simulations (Fig. S4).

The prediction from linear stability theory that mode n=4 should be dominant (over a 2 μm domain) also appears not to hold (Fig. S2b, left panel, red line). The final pattern can consist of a peak or peaks anywhere on the domain though with higher likelihood somewhat away from the boundary (Fig. S4). Note that the dominant mode is not a good descriptor of the pattern in this case as the pattern is generally the sum of many modes, both even and odd (Fig. S2b, left panel, blue lines). In the stochastic reduced model, a peak was mostly situated at one of boundaries, often jumping between them (Fig. S4). In the full system however, the flux balance mechanism evidently shrinks the basin of attraction for the boundary patterns so that they occur less frequently in the deterministic model, while in the stochastic model, the stochastic effects allow the system to jump out of these ‘local minima’ and maintain correct positioning (Fig 1, Fig. S5, Fig. S8).

### Stability of Boundary Modes

To investigate this stability further, we spatially perturbed the patterns obtained by in full deterministic model (over a 2 μm domain) and let them re-stabilise. We did this by cyclically translating the profile of *v*_*m*_ by 0.25 μm. Non-spatial perturbations, in the sense of modifying the relative concentrations of *u*_*m*_, *v*_*m*_ and *u*_*c*_ at each spatial coordinate without mixing between the spatial locations, had little effect. We found that after a boundary focus (n=1) was shifted inwards, it re-stabilised at mid-domain (n=2^-^), whereas a mid-domain focus returned to mid-domain. Thus, the spatial perturbation leads to selective mode transitions consistent with the observed behaviour in stochastic simulations.

### Bifurcation Analysis

To investigate the stability of the mode 2^-^ and 4^-^ patterns and the transition between them, we performed a bifurcation analysis. We treated the length as a bifurcation parameter, increasing or decreasing it in small steps (0.01 μm), starting from either the 2^-^ or 4^-^ pattern. At each step the solution was rescaled to match the domain length and the system solved for 60 min. We then recorded the dominant mode as for the pattern analysis. For measuring the dependence of the one-to-two peaks bifurcation on the diffusion rates, we used a step size of 0.1 μm.

### Parameter Sensitivity

The model has two unconstrained parameters that are important for development of a Turing pattern, namely β and γ. We therefore explored the range over which these parameters result in pattern formation in the stochastic model (the Turing space). We found that patterns were possible at least an order of magnitude above and below our chosen values (the coloured region in Fig. S1c). We also examined the nature of the patterns. We calculated the first 20 wave mode amplitudes of the pattern for different β and γ (20 was found to numerically to sufficient over the parameter range investigated). In Fig. S1c, we have indicated the dominated (non-zero) mode and the fraction of the spectral density associated to it. The dominant mode contributes more to the resultant pattern for the deterministic model in the presence of cytosolic exchange than without (middle row), indicating a pattern more similar to an individual mode. However odd dominant modes, or even dominant modes with positive amplitude, still sometimes occur (as discussed in the text and Fig. 1 and Fig. S2). In the stochastic model, even modes with negative amplitude dominate indicating regular positioning.

### Experiments

Strain AB45 (AB1157 MukB-GFP) (20) was grown in M9 minimal media supplemented with 0.2% glycerol and 0.2% Casamino acids at 30ºC. Fresh media was inoculated from overnight cultures and grown for approximately 4 hrs to an O.D. of 0.1. To stain the nucleoid Sytox® Orange (Thermo Fisher Scientific) (42) was added to an aliquot at a final concentration of 500nM and allowed to grow for a further 20 min. Cells were placed directly (without washing) on a microscopy depression slide prepared with media and 1% agarose. Imaging was performed on a Nikon Ti-E upright microscope with a 100x NA 1.45 phase contrast objective and an Andor Xyla sCMOS camera. During snapshot acquisition, z-stacks of phase contrast and fluorescent images were taken with 200nm spacing (6 slices) and with a 300nm z-offset between the phase contrast and fluorescent (GFP and Sytox Orange) channels. These stacks were then combined into single images with a higher depth of field using the ImageJ Extended Depth of Field plugin. Image segmentation and signal profile extraction was performed using Microbetracker (43). Matlab® was used for the subsequent analysis. Signal profiles were first normalized by area to convert them to intensities (corresponding to protein concentration). To make different cells comparable, we then normalized by the intensity profile by the total intensity in the cell. Cells having the same length, to the nearest two pixels (130nm), were then grouped and their profiles averaged. These profiles were then arranged into an averaged demograph.

## Acknowledgments

We thank David Sherratt for providing strain AB45 and to Remy Colin, Ulrike Endesfelder, Peter Graumann, Suckjoon Jun, Martin Howard and Anja Paulick for discussions and/or comments on the manuscript. This work was supported by the German Federal Ministry of Education and Research and the Max Planck Society in the framework of the research network MaxSynBio.

## Author Contributions

S.M.M. initiated the work, conceived the model and designed and performed simulations and experiments. V.S. contributed to experiment design. Both authors discussed the results and implications. S.M.M wrote the initial draft of the manuscript. Both authors edited subsequent and final versions of the manuscript.

